# Nature based measures increase freshwater biodiversity in agricultural catchments

**DOI:** 10.1101/672915

**Authors:** Penny Williams, Jeremy Biggs, Chris Stoate, John Szczur, Colin Brown, Simon Bonney

## Abstract

This is the first study that describes the effect of adding mitigation measures on the freshwater biodiversity of all waterbody types in agricultural catchments. We measured alpha (site) and gamma (catchment) richness annually over a nine-year period in all the streams, ponds and ditches in three upper-catchments in the English lowlands, and investigated whether freshwater plant biodiversity could be increased by adding: (i) multi-functional ecosystem services measures to intercept pollutants, store water and promote biodiversity, and, (ii) biodiversity-only protection measures. In the absence of measures, all catchments saw a decline in macrophyte richness during the survey (mean species loss of 1% pa, rare species loss of 2% pa). Ponds were a key habitat with a disproportionate influence on catchment trends. Five years after introducing measures, natural colonisation of ecosystem services waterbodies (dammed streams and ditches, runoff ponds, flood storage ponds) largely cancelled-out the background loss of plant species but, importantly, did not restore the loss of rare plants. Adding clean water ponds as a biodiversity-only enhancement measure brought substantial benefits: increasing total-catchment richness by 26%, and the number of rare plant species by 181%. Populations of spatially restricted species also increased. Adding stream debris-dams as a biodiversity measure did not affect plant richness or rarity. The findings suggest that ecosystem services measures could bring some biodiversity benefits to agricultural catchments. However, creating clean-water ponds specifically targeted for biodiversity could hold considerable potential as a tool to help stem, and even reverse, ongoing declines in freshwater plant biodiversity across farming landscapes.

## 1 Introduction

Measures to protect the aquatic environment within farming landscapes currently cost many millions of pounds annually, with expenditure exceeding £470 m per annum in the UK alone (Rayment, 2017). This spend reflects a widespread recognition that agriculture, which makes up about 40% of land cover worldwide and 70% in Britain (Foley *et al.* 2005; Brown *et al.* 2006), plays a major role in modifying, and commonly degrading, freshwater ecosystems and the services that they provide (Moss, 2008; Gordon, *et al.* 2008). In Europe, for example, member states report that nutrient pollution significantly degrades 28% of all surface water bodies classified in the Water Framework Directive (Carvalho et al., 2018).

The mitigation techniques used to protect freshwaters from agriculture-associated impacts are wide-ranging. They include pollutant control measures (e.g. riparian fencing, buffer strips, constructed wetlands, nutrient management plans, minimum tillage), measures to hold back water in catchments, reduce flow, and increase infiltration (e.g. balancing ponds, rural Sustainable Urban Drainage Schemes (SUDS), afforestation), and measures to improve biodiversity and resilience (e.g. debris dams, flow deflectors, river restoration, lake biomanipulation, pond creation and management), (Cuttle et al., 2016, Zhang et al., 2017).

Despite the cost and effort involved in implementing these measures, meta-analyses suggest they are variable in their effectiveness and improvements in physico-chemical and ecological measures are often considerably less than anticipated. This is certainly true for freshwater biodiversity where few studies have proven that measures bring significant biodiversity gains (Bernhardt and Palmer, 2011; Louhi et al., 2011, Harris and Heathwaite, 2012; Robertson et al., 2018). Our limited understanding of the benefits of agricultural measures on freshwater biodiversity is compounded by an almost universal tendency for studies to evaluate success based on a very partial element of the catchment: mainly rivers and streams. We now know that small waterbodies including ponds, springs, headwaters and ditches, typically support a high proportion of the freshwater biodiversity within agricultural catchments (Williams et al., 2004; Céréghino, 2007, Davies et al., 2008, Bubíková and Hrivnák, 2018). Systematically ignoring these habitats when assessing the benefits of mitigation measures can easily lead to a perception bias. For example, because there is a considerable overlap between the biota of standing and running waters (Biggs et al., 2017), species gains reported in a river network can be trivial if these taxa are already widely present in other catchment waterbodies. Likewise, simply relying on data from streams and rivers can miss broader trends that are profoundly affecting freshwater biodiversity across the catchment as a whole.

In this paper we look at the impact of introducing a range of mitigation measures on higher plant biodiversity in all waterbodies within a typical agricultural area of the English Midlands. The mitigation measures we applied included common ecosystem services measures, primarily intended to slow flows, intercept polluted water and sediment and store flood water, including bunded (i.e. dammed) streams and ditches, interception ponds and flood storage ponds. We also added two simple habitat creation measures specifically intended to bring long-term biodiversity benefits: clean water ponds and stream debris dams. Clean water ponds are off-line waterbodies (not connected to streams or ditches) located in parts of the landscape where, as far as possible, they fill with unpolluted surface-water or groundwater. Recent evidence suggests that these waterbodies can rapidly become species-rich and retain their value for many years (Williams et al., 2008, 2010; Oertli, 2018). Debris dams are widely applied in river restoration as features intended to increase habitat (and therefore by implication, biotic) diversity (Roni et al., 2014). Because our measures not only target ecosystem services but address wider societal challenges such as maintaining biodiversity in an era of accelerating extinction and loss (Dullinger et al., 2013; IPBES 2019), we have termed the combination of these interventions ‘nature based measures’. This references the broader term ‘nature-based solutions’ used by IUCN and other organisations such as the European Commission when referring to ‘actions [that] protect, sustainably manage and restore natural or modified ecosystems’ (Cohen-Shacham et.al. 2016).

To assess the effect of our nature based measures we undertook site and whole-catchment studies in all the freshwaters present in the study area(streams, ponds and ditches). The landscapes have no waterbodies large enough to be described as rivers or lakes (Brown et al., 2006). We used wetland plant attributes to measure biodiversity because plant data can be collected relatively quickly and easily. This enabled us to study real species loss and gain in our catchments through a census survey of all plant species in all waterbodies, rather than relying on calculated estimates of gamma richness, which is more typical of catchment studies (Williams et. al., 2004, Davies et. al., 2008). In addition, temporal and spatial trends in wetland plants have been shown to correlate positively with trends in other biotic groups, particularly aquatic macroinvertebrates, making wetland plants a broadly representative group (Williams et al., 2004; Zelnik et al., 2018; Law et al., in press).

Our study was based on a Before and After design with a three-year baseline and, to date, five years of post-intervention monitoring. The work forms part of the Water Friendly Farming initiative: a long-term research and demonstration project to investigate the effectiveness of landscape-wide mitigation measures intended to reduce the impact of rural land use on water, whilst maintaining profitable farming (Biggs et al., 2014).

## 2 Methods

### 2.1 Study area

The Water Friendly Farming study area lies within three sub-catchments of the River Welland and the River Soar in Leicestershire, England. Each catchment is around 10 km^2^ in area (Table 1, Figure 1). The catchments are directly adjacent and have very similar geologies, topographies and land uses. This is a region of low rolling hills (95-221 m OD) with high ground areas (including most headwaters) dominated by Pleictocene fluvio-glacial sands, gravels and clays. Valley sides are predominantly Middle and Lower Jurassic mudstones and siltstones with beds of ferruginous limestone. In valley bottoms the Jurassic strata are overlain by recent deposits of alluvium or colluvium. Cultivated land falls into two of the most extensive of Britain’s agricultural land classes: Defra Land Class 4 eutrophic tills, and Land Class 6, pre-quaternary clay, which together make up 35% of the cultivated land in Great Britain (Brown et al., 2006). Agriculture in all catchments is mixed farming: divided between arable land mainly under oilseed rape and winter wheat with additional field beans or oats, and grassland used to pasture beef cattle and sheep or cut for hay or silage. Table 1 shows the proportion of land cover types in each catchment. Maps showing the distribution of land cover and waterbody types are provided as supplementary information.

**Figure 1.**
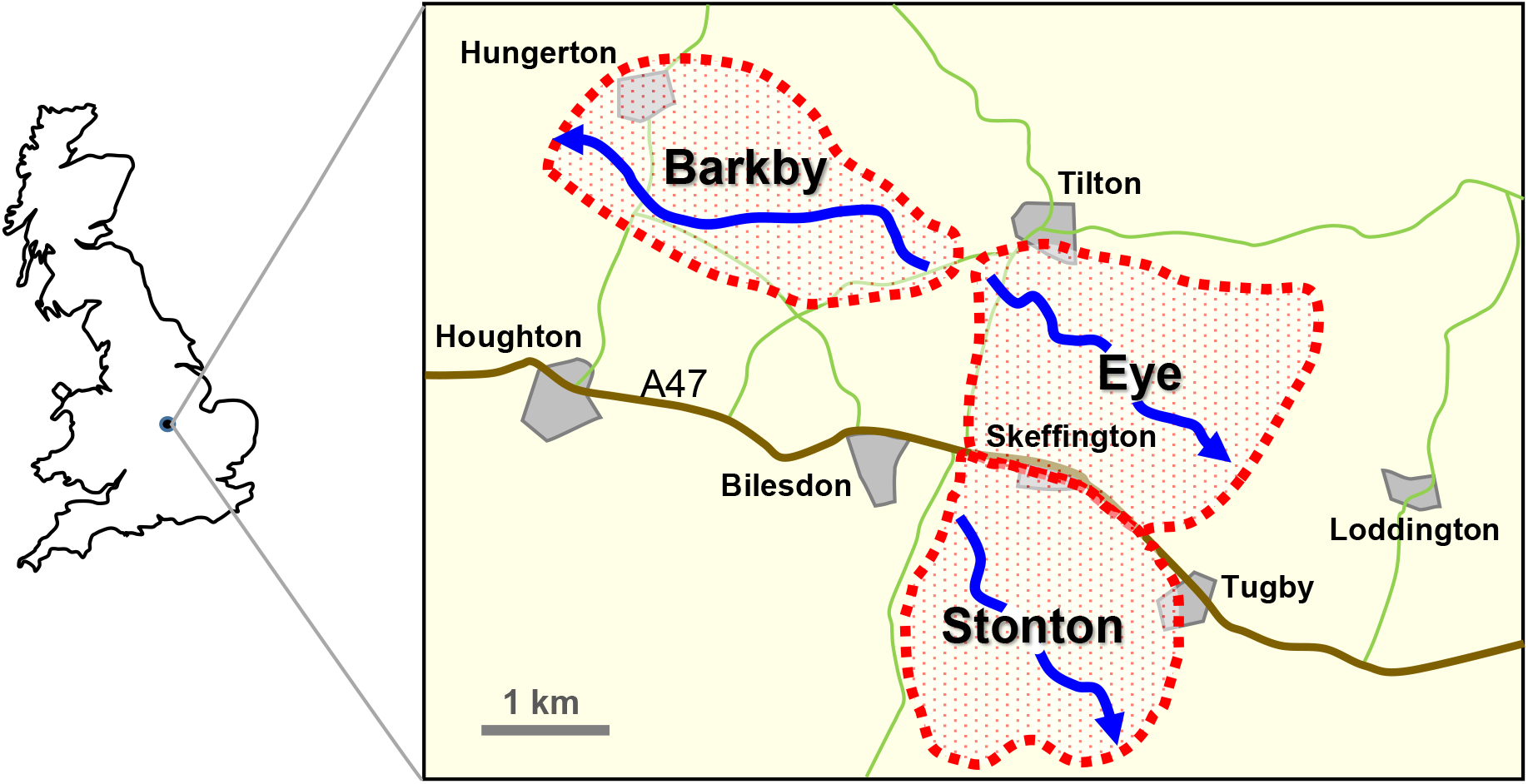
Location of the three survey catchments

The study area has three main waterbody types based on the definitions given in Williams et al. (2004): streams, ditches and ponds. Streams in all catchments are relatively small with a maximum width of c.3m. At the time of the survey the lower reaches of all streams were typically shaded by riparian trees with their margins supporting shade tollerant plants such as *Epilobium hirsutum, Solanum dulcamara, Angelica sylvestris* and *Filipendula ulmaria.* In the upper reaches of all catchments some stream lengths were more open and bordered by fenced or unfenced pasture often with the channel supporting *Veronica beccabunga*, *Juncus effusus* and *Glyceria notata.* Ditches were typically narrow (0.5-1.5m width) and borded by a hedge on one side with *Epilobium hirsutum,* and *Solanum dulcamara* the most common wetland plants. Most ditches, and the headwaters of some streams were seasonal for part of the year. Ponds showed considerable between-waterbody differences in terms of their shading and seasonality. In pastoral areas most ponds were fenced. In addition to supporting the plant species that were widespread in streams and ditches, ponds commonly supported grasses and rushes such as *Juncus inflexus*, *Deschampsia cespitosa*, *Glyceria fluitans* and *Sparganium erectum*. They were also the only habitat to consistently support floating and submerged leaved plants, with the most widespread taxa: *Ceratophyllum demersum*, *Potamogeton berchtoldii* and *Lemna minor*.

### 2.2 Experimental set-up

The study is based on a Before and After design with two experimental catchments centred on the Eye (52° 37’ 34.4”N 00° 53’ 12.4”W) and the Stonton (52° 36’ 08.0”N 00° 54’ 32.8”W). A third catchment, the Barkby (52° 38’ 48.9”N 00° 54’ 28.0”W), provides an additional control site where no measures were added (Figure 1). Survey waterbodies were identified using a combination of Ordnance Survey 1:1 250 scale maps, landowner information and field-walks. Nature based mitigation measures were introduced in 2013 and the early months of 2014, with the exception of debris dams (most introduced in 2015). All measures were left to colonise naturally, with no wetland plants added to the waterbodies or their banks.

Two types of measures were added (Table 1):

i. Ecosystem services measures introduced to both the Eye and Stonton catchments. These were designed to have a multifunctional role including pollution reduction, flood peak attenuation, groundwater recharge and biodiversity protection. They included (a) earth-bunded (i.e. dammed) streams and ditches mainly used to hold-back water and trap sediment and (b) two types of interception ponds: run-off ponds that intercept arable field drains and flood-storage ponds filled by streams or ditches during periods of high water flow.
ii. biodiversity-focussed measures, which were added to only one of the experimental catchments (Stonton). Two measures were added: clean water ponds and stream debris dams, both specifically designed to increase catchment species richness. The third catchment (Barkby) remained as a control. No mitigation measures were installed here although, as in the Eye and Stonton, this catchment had a normal level of agri-environment scheme protection (e.g. cross compliance buffer strips, Defra 2018).

### 2.3 Sampling strategy

All catchments were monitored for three years prior to the addition of nature based protection measures (2010-2012). Ecological surveys were not undertaken during 2013 whilst measures were put in place. Post-mitigation monitoring was undertaken annually during the five-year period from 2014 to 2018.

Wetland plant (i.e. aquatic macrophyte) data were collected annually from waterbodies. ‘Wetland plants’ were defined as the plants listed in Freshwater Habitats Trust (2015) Wetland Plants Recording Form, which comprises a standard list of ca. 300 water-associated higher plants divided into three categories: submerged, floating-leaved and emergent plants. Plant species and their percentage abundance were recorded while walking and wading the margins and shallow water areas of the waterbody. For sites with deeper water, submerged aquatic plants were surveyed using a grapnel thrown from the bank. To ensure consistency a single surveyor undertook all surveys. The main botanical survey was undertaken in August each year, however, an additional check was made for early-growth plant species in May, when aquatic invertebrate surveys were undertaken. This enabled Batrachian *Ranunculus* species to be identified whilst in flower.

Site (alpha) richness data were collected from randomly selected locations that were revisited each year (termed here ‘standard samples’). Site selection was stratified by catchment and waterbody type. Twenty standard sample sites were surveyed from each waterbody type in each catchment. This gave a total of 360 sample sites per year prior to the introduction of measures, and around 420 thereafter. River and ditch sections were selected by dividing the network into 100 m lengths and randomly selecting 20 lengths for survey. To ensure that ecological data gathered from different waterbody types could be directly compared, the sampling was area-limited with data from each site collected from a 75 m^2^ area of the waterbody based on the method described in Williams et al. (2004). Although this area-based method enabled waterbodies with widely differing dimensions and characteristics to be compared, small waterbodies less than 75 m^2^ are by definition, excluded from the survey. To avoid completely omitting smaller habitats, where appropriate, closely adjacent pools were aggregated to give a 75 m^2^ total area. This included tree-fall pools in a wooded fen, and small ecosystem services bunded-ditch pools in a series of two or three features. Debris dams were assessed by surveying a 75m^2^ area of stream centred on the dam.

Gamma richness data were collected from a census survey of all ponds and ditches in each year. Streams were an extensive habitat type that could not be fully surveyed annually. Our original aim was to use rarefaction curves (Colwell et al. 2012), to calculate gamma richness from a dataset combining standard samples and an additional 10 randomly selected stream samples. In practice the rarefaction curves gave variable and intuitively unlikely results. To investigate this, all streams were fully surveyed in 2018. Comparison of our stream census data and rarefaction-predicted results showed that rarefaction curves over estimated true gamma richness in 2018 by 17%-68% depending on the algorithm used and catchment modelled. Simple summing of the standard and random alpha richness data gave a result that was a close match to the true gamma of our relatively homogeneous streams: varying between zero and one species per catchment below true gamma. In the following analysis we have therefore used the summed stream alpha survey data to represent true gamma for all years.

**Table 1.**
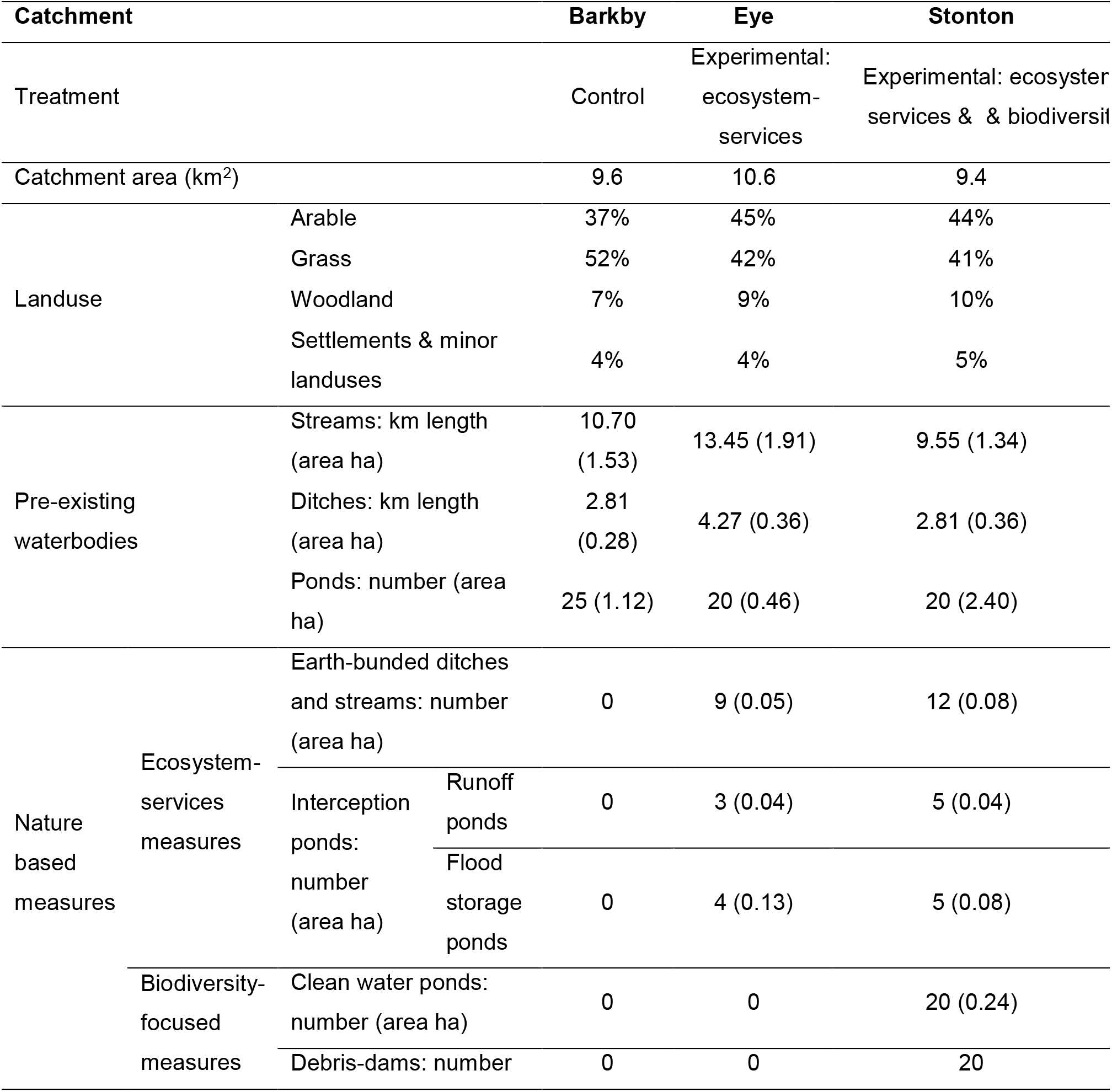
Physical characteristics of the three survey catchments and nature based features added by the project. The table shows the length and number of waterbodies, with their total area in parentheses

### 2.4 Analytical methods

#### 2.4.1 Waterbody dimensions

The dimensions of pre-existing waterbodies were calculated from 1:500 scale map data and ground-truthed in the field. The area of new ecosystem services and biodiversity features was measured as dug. The proximity of standing waters was calculated as a nearest neighbour analysis on ArcGIS Pro 2.3.

#### 2.4.2 Biodiversity

Wetland plant biodiversity was assessed on the basis of species richness and species rarity. Alpha richness was measured as the number of species, or distinctive taxa, recorded at each survey site. Alpha rarity was the number of regionally or nationally rare plant species recorded in any of the following categories: (i) regionally rare species - recorded from fewer than 15% of 1 km grid squares in a 100 km^2^ grid square centred on the project area (BSBI, 2019), (ii) Nationally Scarce species - recorded from 16 to 100 10×10 km grid squares in Britain (JNCC, 2019), (iii) species that are Red Listed in England or at UK level - based on the IUCN categories: Near Threatened, Vulnerable, Endangered or Critically Endangered (Stroh et al., 2014). We also calculated the rarity of species within our catchments using two measures: (i) the number of species with restricted distribution in the three catchments, measured as species recorded from only one or two sites, termed ‘uniques’ and ‘doubles’ respectively (c.f. Gotelli and Colwell, 2011), (ii) the number of restricted species with small populations, measured as species with a total aerial coverage of <5m^2^.

Gamma richness and rarity were calculated as the total number of species and rare species recorded in each waterbody types and catchment area. To enable comparison between our gamma results and other studies we recalculated gamma richness data from Hassall (2012, Table 6) based on the more restricted wetland-only plant species list used in the current study (see Supplementary information).

#### 2.4.3 Statistical methods

Statistical differences between the species richness of waterbody types and catchments were tested using two and three way between-subjects ANOVAs, using square-root transformed data. Non-parametric tests (2-tailed Mann-Whitney U and Friedman tests) were used to analyse plant rarity data where there were a high proportion of zeros. Simple linear regression was used to investigate changes in richness and rarity over time. Analyses were run on IBM SPSS Statistics version 2015.

## 3 Results

### 3.1 Baseline wetland plant results for the three catchments

Underlying trends in alpha (site) and gamma (catchment) plant richness and rarity were calculated excluding new ponds and other waterbodies created or modified after 2013 as part of project measures. The results therefore indicate the background biodiversity trends in the absence of the direct effect of the project’s physical habitat creation work. Species lists for each waterbody type and catchment are provided as supplementary information.

#### 3.1.1 Background alpha richness and rarity trends

Plant richness differed among waterbody types (*F*(2,1368) = 165.65, p < 0.001) and was greatest in ponds (Mean=8.93 species, SE =0.20) followed by streams (M=5.58, SE=0.18) and then ditches (M=4.28 SE=0.13). Ponds were the only habitat type to support submerged and floating-leave plant species consistently (Figure 2). There were differences in plant richness among the three catchments (*F(*2,1368) = 20.960, *p*<0.001). Alpha richness was significantly greater in the Barkby than in the Eye and Stonton catchments (M=7.19 SE=0.21), (*p*<0.001 both comparisons). Differences between the Eye and Stonton were not significant (M=6.16 SE=0.19; M=5.44 SE=0.18 respectively), (*p*=0.07). There was no statistical evidence for a temporal trend in alpha plant richness across the survey period, with no significant main effect for year or for the threeway interaction and between-subject effects for year, waterbody type and catchment (Figure 2).

**Figure 2.**
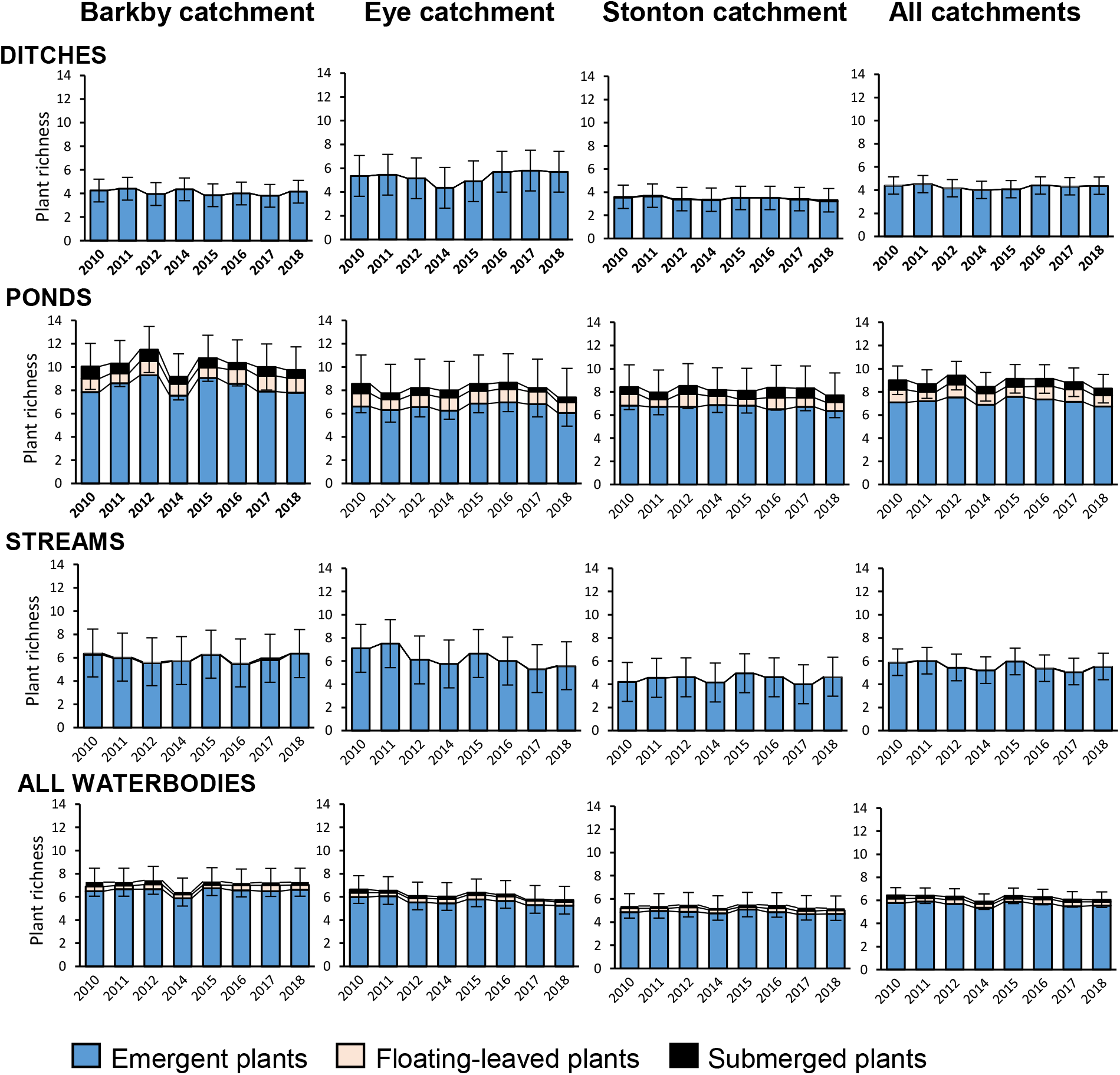
Alpha richness in all waterbody types and catchments: showing the mean number of wetland plant species recorded from an annual survey of 75m^2^ plots from 20 ditches, ponds and streams in each of three catchments. No data were collected in 2013 when measures were being introduced. The graphs do *not* include new waterbodies or features added after 2014, and hence show underlying trends in the absence of nature-based measures. Error bars show 95% CLs for total mean richness.

The majority of rare plant species were restricted to ponds (mean of 0.44 species per site). The occurrence of rare plants in streams and ditches was an order of magnitude lower (both 0.03 rare species per site). The high proportion of zero values in the dataset precludes tests of statistical significance. The number of rare plant species did not differ significantly between any of the three catchments (Mann-Whitney U test). There was no evidence of temporal trends in the rare species data for any catchment (Friedman test of differences among repeated measures).

**Figure 3.**
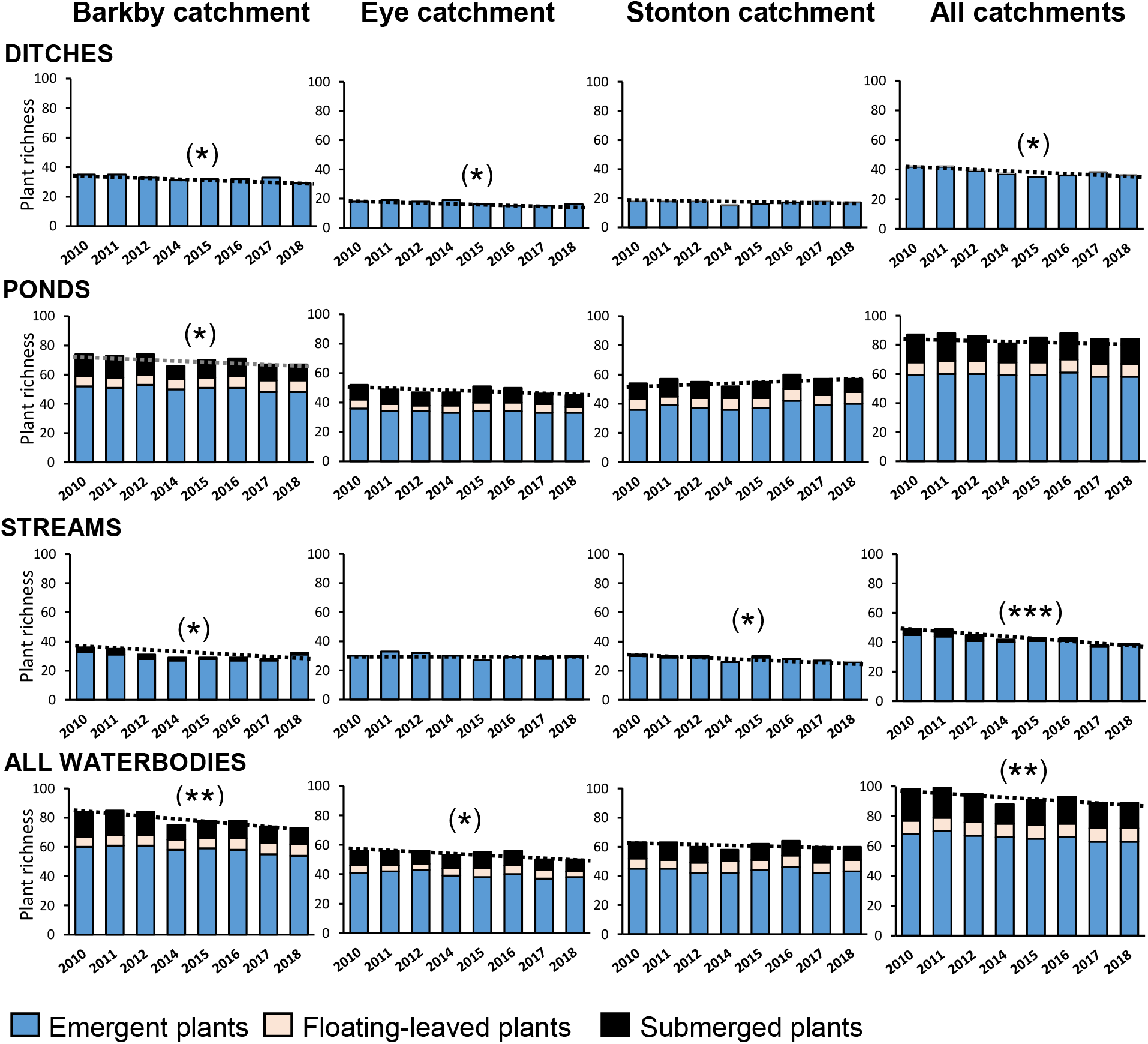
Plant gamma richness for all ditches, ponds and streams in each of three catchments, shown with a line of best fit. No data were collected in 2013 when measures were being implemented. The graphs do *not* include new waterbodies or features added after 2014, and hence show underlying trends in the absence of nature based measures. Dotted lines show the simple linear regression for total plant richness in each waterbody and catchment. Asterisks denote the statistical significance of the regression equation: p-value <0.05 (**), < 0.01 (**), < 0.001 (***).

#### 3.1.2 Gamma richness and rarity trends

In the absence of measures, a total of 106 wetland plant species were recorded from census surveys of the three catchments during the survey, with a mean of 92.8 species (range 89-99) in any one year. The Barkby control catchment supported the greatest number of species per year (78.9 species, 85% of total gamma for all three catchments), followed by the Stonton (61.3 species, 66% of total gamma) and Eye (54.0 species, 59% of total gamma). Gamma richness differences between the waterbody-types broadly concurred with the alpha richness trends. Ponds were much the richest waterbody type in all three catchments in all years, supporting an average of 85.5 species per year. Streams and ditches each supported around half this (43.5 and 38.9 species, respectively).

Analysis of temporal trends shows that, in the absence of measures, the number of plant species present in all waterbodies and catchments declined by 10% during the nine year survey period, (F(1,6)=13.79, *p*<0.01) with an adjusted *R^2^* of 0.65. (Figure 3). This represents a loss of 1.1 wetland plant species per annum across the combined area of the three catchments.Of the individual catchments, the Barkby catchment, which was initially the most species-rich, saw the greatest decline in plant species with a 14.0% loss through the survey period: and average of 1.4 species per year F(1,6)= 29.74, *p*<0.01) with an adjusted *R^2^* of 0.80. The Eye catchment declined by 10% F(1,6)= 7.99, *p*<0.05) with an adjusted *R^2^* of 0.50, 0.6 species per year. Declines in the Stonton catchment were smaller (2%) and not statistically significant F(1,6)= 0.34, *p*=0.58) with an adjusted *R^2^* of −0.10.

Amongst individual waterbody types, streams declined at a rate of 1.2 species per annum (F(1,6)= 43.77, *p*<0.001) with an adjusted *R^2^* of 0.86. Loss in ditches was lower (0.7 species per annum, F(1,6)= 9.76, *p*<0.05, with an adjusted *R^2^* of 0.57). Pond richness did not decline significantly across the three catchments, (F(1,6)= 1.55, *p*=0.26) with an adjusted *R^2^* of 0.07, although there was a significant decline in the richness of Barkby catchment ponds alone (*p*<0.05). Across the dataset, pond gamma richness patterns in the Stonton catchment were anomalous as the only catchment waterbody type where, in the absence of measures, more species were present at the end of the survey than at the start (Figure 3). Examination of the data suggests that at least part of this increase is likely to have been due to an unexpected indirect effect of the project’s habitat creation after 2014: with new-to-the-catchment species, like the submerged aquatic *Potamogeton pusillus* and marginal plant *Equisetum palustre,* that rapidly colonised the new clean-water ponds, subsequently moving out to colonise adjacent pre-existing ponds.

In total, 18 plant species that were either nationally or regionally rare were recorded from the baseline waterbodies in the three catchments across all years; 16% of the total flora. Ponds supported by far the greatest proportion of these rare species (89% across all catchments and survey years). Streams and ditches each supported 17% of the rare species pool.

All catchments showed a tendency towards a background loss of rare plant species during the survey (Figure 4). Simple regression of total gamma rarity for all waterbodies and catchments combined gives a decline of 22% (0.4 species pa), (F(1,6)= 8.09, *p*<0.05) with an adjusted *R^2^* of 0.50. Rare plant loss was greatest in the Barkby where there was a loss of 0.5 rare species per annum (34% decline in the Barkby’s rare plant species during the sampling period), (F(1,6)= 13.22, *p*<0.05) with an adjusted *R^2^* of 0.64. Rare species loss was also high in the Eye catchment: 0.3 species per annum (38%), (F(1,6)= 10.46, *p*<0.05) with an adjusted *R^2^* of 0.58. The Stonton catchment showed a small net decline in rare species, but the regression relationship is not significant (F(1,6)= 0.58, *p*=0.48) with an adjusted *R^2^* of −0.06. In all catchments, loss of rare plant species was predominantly (63%-75%) due to the loss of submerged aquatic plant species from the catchments’ ponds.

#### 3.1.3 Unique and restricted species in the catchments

Combining data from all catchments and all years shows that close to half (48%) of the plant species recorded from baseline waterbodies were found in only one of the three waterbody types. Of these species, 83% were restricted to ponds, 13% to streams and 4% to ditches. This pattern was more striking for rare species where 89% of the baseline plant species were restricted to a single waterbody type, and 88% of these were unique to ponds. Two species were unique to other waterbody types, but both of these were lost early on in the survey due to adverse management practices.

Many plant species had a highly localised distribution and small populations. In the final year of the survey 34% of taxa were found only as uniques or doubles (i.e. recorded from one or two sites respectively) across the three catchments. Of these localised taxa, 90% were found in ponds, and 83% only in ponds; 10% were recorded only in streams and 3% only in ditches. Most uniques and doubles also occurred at low abundance, with 70% occupying a total area of less than 5m^2^ across all catchments. The majority (86%) of these were pond-only species. Combining these data shows that 23% of all remaining plant species across the three catchments were both present at very few sites and occurred in low abundance within those sites. 86% of these highly restricted plant species were unique to ponds.

**Table 2.** Mann-whitney U-test results comparing the alpha plant richness of nature based measures and pre-existing waterbodies. Statistically significant results in bold

|  |  | Bunded | Interception | Clean water | Pre-existing | Pre-existing | Pre-existing |
| --- | --- | --- | --- | --- | --- | --- | --- |
|  |  | ditches & streams | ponds | ponds | ponds | streams | ditches |
| Debris dams | U | 91.500 | 34.500 | 0.000 | 86.000 | 247.000 | 305.000 |
|  | Z | -3.124 | -4.281 | -5.493 | -4.944 | -2.415 | -1.505 |
|  | p | **.002** | **<.001** | **<.001** | **<.001** | **.016** | .132 |
| Bunded ditches & streams | U |  | 89.500 | 5.500 | 227.000 | 394.500 | 291.500 |
|  | Z |  | -2.822 | -5.427 | -2.946 | -0.390 | -1.970 |
|  | p |  | **.004** | **<.001** | **.003** | .697 | **.049** |
| Interception ponds | U |  |  | 32.000 | 358.000 | 198.000 | 118.500 |
|  | Z |  |  | -4.438 | -0.034 | -2.735 | -4.082 |
|  | p |  |  | **<.001** | .973 | **.006** | **<.001** |
| Clean water ponds | U |  |  |  | 85.500 | 49.000 | 7.500 |
|  | Z |  |  |  | -5.091 | -5.645 | -6.282 |
|  | p |  |  |  | **<.001** | **<.001** | **<.001** |

### 3.2 Effect of adding nature based measures

#### 3.2.1 Physical effect of adding measures

Adding ecosystem services measures to both the Eye and Stonton catchments increased the area of freshwater habitat by an average of 0.2 ha per catchment (Table 1). Adding biodiversity-only measures to the Stonton catchment alone added an extra 0.24 ha. Proportionally, the nature based measures increased the area of standing waters present in the Eye and Stonton by an average of 33%, and the total area of all freshwater habitats in these catchments by an average of 9%.

Nearest neighbour analysis showed that, in the Eye catchment, adding ecosystem services measures reduced the between-waterbody distance of standing waters by 29% from an average of 336 m to 240 m. In the Stonton, where clean water ponds were also added, there was a 64% reduction in distance from 255 m to 92 m between standing waterbodies.

#### 3.2.2 Alpha richness and rarity of measures waterbodies

In the final survey year, five years after their creation, the mean alpha richness of the new clean water ponds was significantly greater than other nature based waterbody types created (p <0.001 for all analyses, Figure 5a). Plant richness associated with debris dams was universally low, and significantly less than for other measures (Table 2). Amongst the ecosytem services measures, interception ponds were significantly richer than bunded ditches and streams (p <0.01). The alpha richness of the new clean water ponds was also significantly greater than the richness of all pre-existing waterbody types in the two experimental catchments (Table 2). The richness of interception ponds was similar to pre-existing ponds. Bunded ditches and streams had a similar mean richness to pre-existing streams, and were marginally richer than pre-existing ditches (p=0.049), (Figure 5).

Stream richness adjacent to debris dams was significantly lower than was typical of pre-existing streams (p<0.05), and was more similar to the richness of ditches. It is likely that this is because debris dams were typically added to smaller streams, most of which were also heavily shaded.

**Figure 4.**
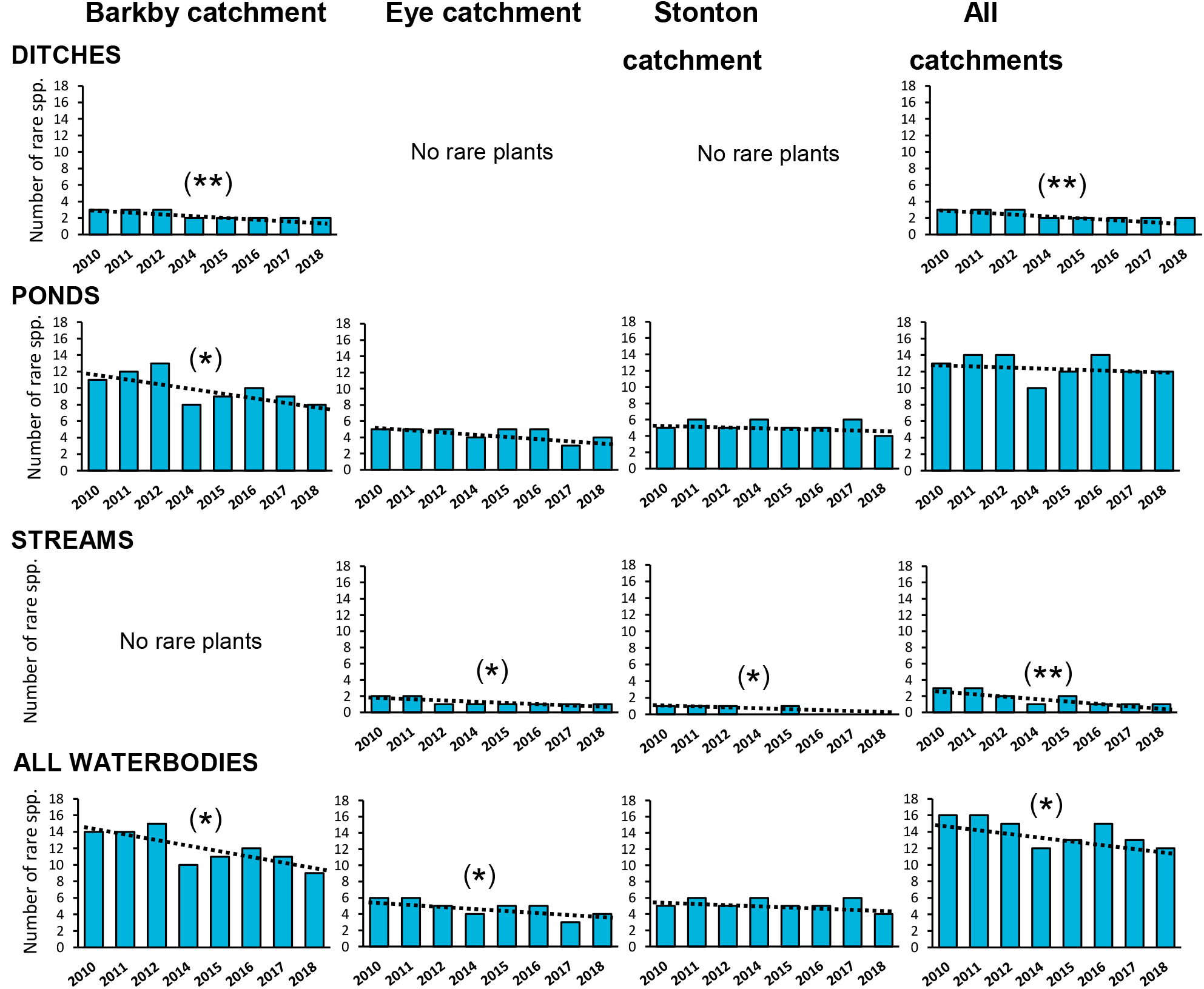
Plant gamma rarity for all ditches, ponds and streams in each of three catchments, shown with a line of best fit. No data were collected in 2013 when measures were being implemented. The graphs do *not* include new waterbodies or features added after 2014, and hence show underlying trends in the absence of nature based measures. Dotted lines show the simple linear regression for total plant rarity in each waterbody and catchment. Asterisks denote the statistical significance of the regression equation: p-value <0.05 (**), < 0.01 (**), < 0.001 (***).

This is supported by comparison between debris dams sections and the nearest unaffected stream lengths which shows that their average richness was almost identical (debris dams 2.6 species; adjacent streams 2.7 species).

Of the four main types of nature based measures, only clean water ponds and interception ponds supported rare species. At five years old, the mean alpha rarity of clean water ponds exceeded that of other nature based and pre-existing waterbody types and was around double that of pre-existing ponds (Figure 5c). The high proportion of zeros values in the data set precludes tests of statistical significance.

#### 3.2.3 Gamma richness and rarity of measures waterbodies

Amongst the introduced measures, clean water ponds supported the greatest gamma richness, followed by interception ponds, bunded watercourses and debris dams (Figure 5b). However, the gamma richness of both pre-existing ponds and streams exceeded the total richness of any of the measures. Gamma rarity showed a different trend (Figure 5d) with the new clean water ponds supporting the greatest number of rare species, followed by the pre-existing ponds.

**Figure 5.**
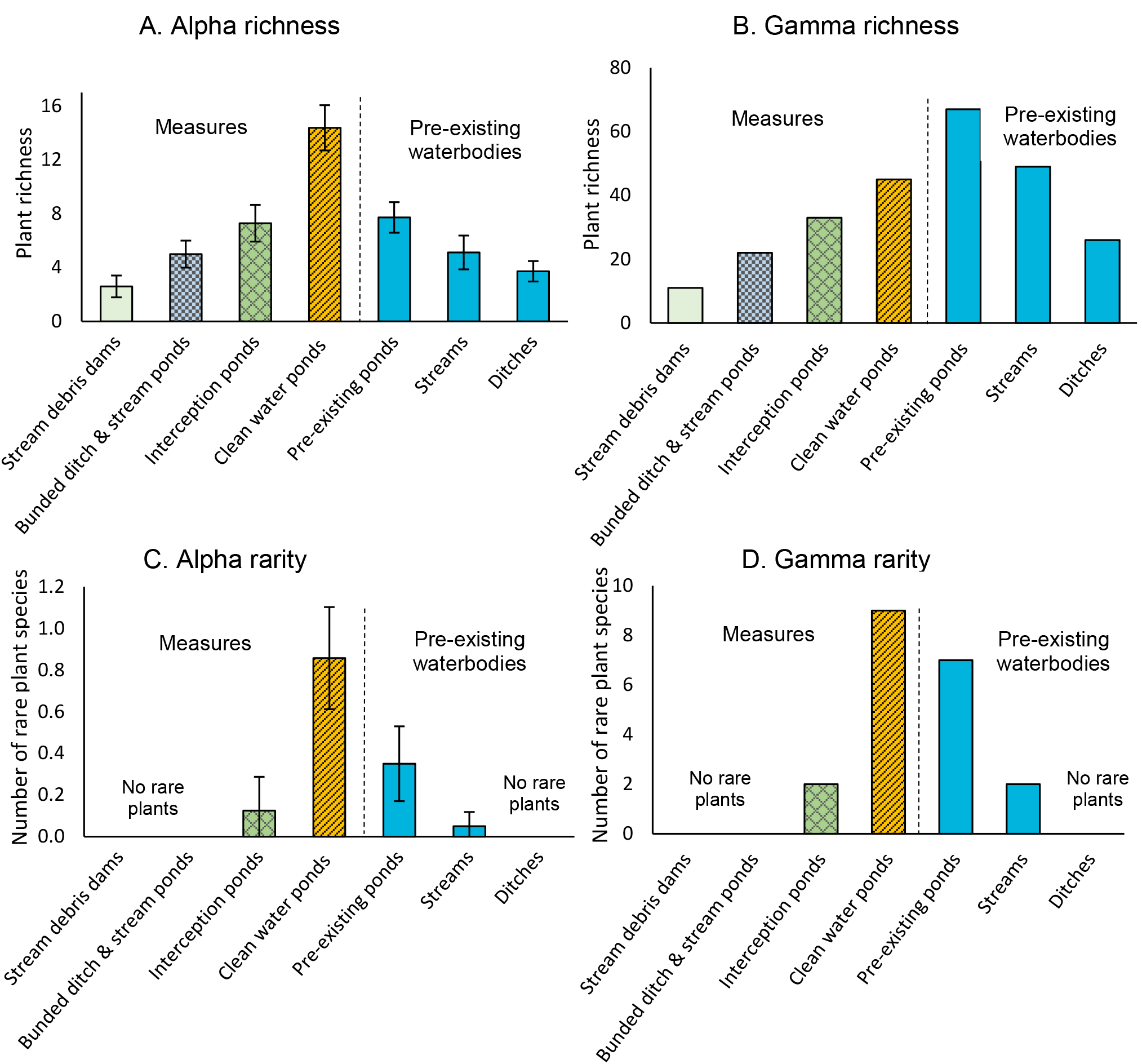
Alpha and gamma richness and rarity of measures and pre-existing waterbodies in the Eye and Stonton catchments in 2018, when the measures were five years old. Error bars for alpha richness and rarity show 95% CLs.

Interception ponds supported few rare species, and they were absent from bunded watercourses and debris dams.

#### 3.2.4 Catchment scale effects from introducing measures

The addition of nature based measures had notable catchment-scale effects in both the Eye and Stonton catchments (Figures 6, 7).

In the last year of the study (2018), when the measures were five years old, new-to-the-catchment plant species in ecosystem services waterbodies increased the total wetland plant richness of the Eye catchment by 14%, from 50 species in all pre-existing waterbodies, to 57 species after addition of the measures (Figure 6a). Simple regression shows that this converted a statistically significant loss of species in pre-existing waterbodies to a small, non-significant gain in the Eye catchment (F(1,6)= 1.44, *p*=0.28) with an adjusted *R^2^* of 0.06. In the Stonton catchment, ecosystem services measures increased species richness by 7%, (Figure 6b), increasing the small non-significant downward trend in richness in pre-existing waterbodies to a small non-significant upward trend (F(1,6)= 1.88, *p*=0.22) with an adjusted *R^2^* of 0.11.

**Figure 6.**
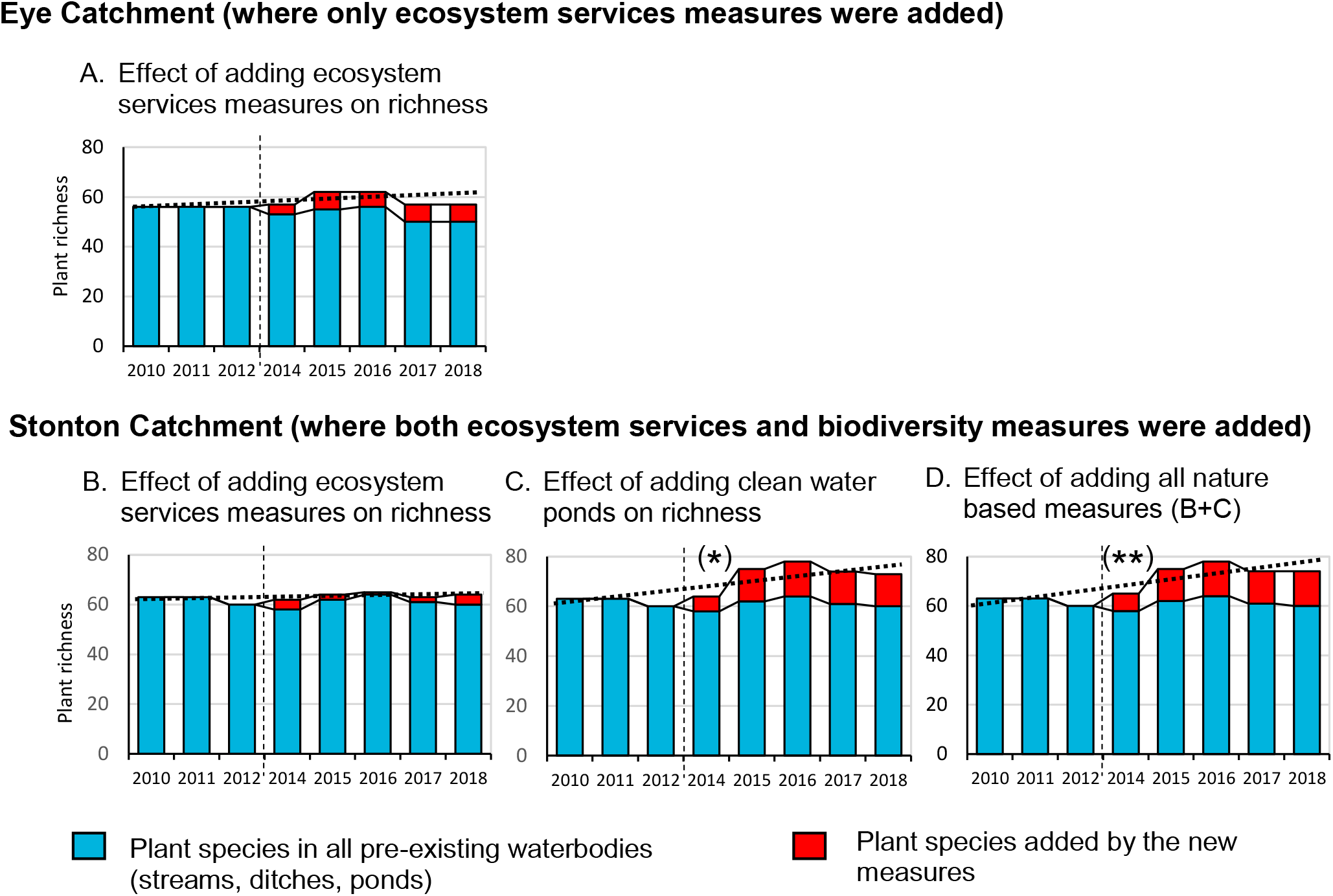
Change in gamma richness as a result of adding nature based measures to the experimental catchments. Ecosystem-services measures were added to the Eye Catchment. The Stonton Catchment received both ecosystem-services measures and clean water ponds. Dashed vertical lines separate the pre- and post-measure phases. Dotted lines show the simple linear regression for total plant richness. Asterisks denote the statistical significance of the regression equation: p-value <0.05 (**), < 0.01 (**), < 0.001 (***).

The clean water ponds contributed more substantially to the Stonton catchment. In 2018, the five-year old clean water ponds added 13 new wetland plant species; a 22% increase from 60 species in all pre-existing waterbodies, to 73 species with the clean water ponds (Figure 6c). Simple linear regression shows that this changed the small non-significant trend towards loss of species in the Stonton catchment to a significant 26% gain (F(1,6)= 12.32, *p*<0.05) with an adjusted *R^2^* of 0.62.

Few plant species were unique to the ecosystem services ponds and debris dams added no new plant species (Figure 6d). Hence the total richness of all nature based measures in the Stonton catchment was similar to the richness added by the clean water ponds alone. The combined effect of adding all nature based measures was a statistically significant increase of 27% based on simple linear regression of data for all years (F(1,6)= 15.12, *p*<0.01) with an adjusted *R^2^* of 0.67.

Both the ecosystem services and clean water ponds supported rare species that were new to, or had recently become extinct from, their catchments (Figure 7). However, in the new ecosystem services waterbodies regionally uncommon species such as the submerged aquatic *Potamogeton pusillus* were transitory colonisers, and did not persist after the features were 1-3 years old (Figures 7a,b). The clean water ponds made a substantial and lasting contribution to catchment rarity. In 2018, the final year of the survey, pre-existing waterbodies in the Stonton catchment supported only four rare species. Adding the clean water ponds tripled this to 12 species. Linear regression for the survey period as a whole, shows that their addition changed the small non-significant trend towards rare species loss of species in the Stonton catchment to a significant gain of 181% (F(1,6)= 35.94, *p*=0.001) with an adjusted *R^2^* of 0.83 (Figure 7c). New rare species that colonised the clean water ponds included the Near Threatened *Triglochin palustris*, together with species such as *Isolepis setacea*, *Juncus subnodulosus* and *Hippuris vulgaris* that are both rare in the region and increasingly uncommon in lowland England.

**Figure 7.**
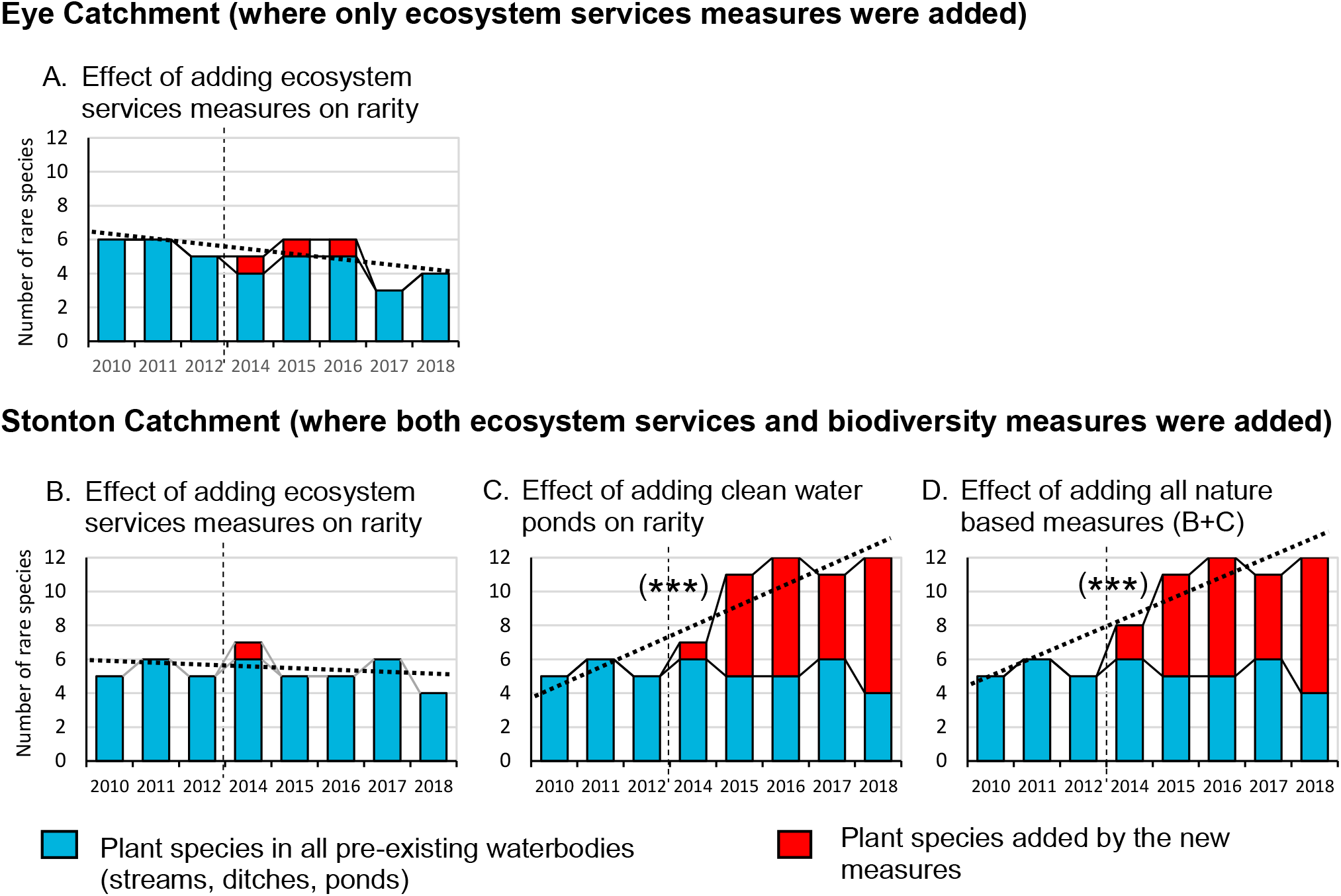
Change in gamma rarity as a result of adding nature based measures to the experimental catchments. Ecosystem-services measures were added to the Eye Catchment. The Stonton Catchment received both ecosystem-services measures and clean water ponds. Dashed vertical lines separate the pre- and post-measure phases. Dotted lines show the simple linear regression for total plant rarity. Asterisks denote the statistical significance of the regression equation: p-value <0.05 (**), < 0.01 (**), < 0.001 (***).

##### The effect of adding nature based measures on catchment rarity

On average, 18.5% of species that had been recorded in the pre-existing Eye and Stonton waterbodies (2010-2017), were no longer recorded in these catchments in 2018. Of these ‘lost’ species 15% were present in the Ecosystem Services waterbodies. In the Stonton catchment where clean water ponds were also created, 31% of ‘lost’ species were present in these ponds. No ‘lost’ species were retained as a result of creating the debris dams.

In the final year of the study both types of nature based measures also created habitats for populations of plants that were otherwise highly restricted in their catchments. In total, the clean water ponds provided 34 new sites for six (22%) of the plant species that were otherwise present only as uniques or doubles in the Stonton catchment. Ecosystem services ponds provided fewer opportunities to secure new populations of vulnerable species, with an average of seven new sites for 3.5 species (12%) of uniques or doubles present in the Eye and Stonton catchment.

## 4 Discussion

### 4.1 Trends in catchment aquatic biodiversity

This study appears to be the first to look at catchment-level temporal trends in wetland plant richness across the full range of waterbody types present in a typical agricultural landscape. The results showed that, in the absence of measures, there was a systematic decline in plant richness over the nine-year survey period, with an annual loss of 1% of plant species, and 2% loss of rare species across the 30 km^2^ survey area as a whole. The limited availability of comparable data makes it difficult to know how typical these trends are of wetland plant assemblages in other lowland agricultural catchments. The main exception is Gołdyn (2010) who undertook a 30 year re-survey of plant richness across a range of agricultural waterbodies in West Poland. She found an increase in gamma richness between 1976 and 2007, although this was mainly due to colonisation by alien and ruderal species, and the number of rare and threatened plants declined. Of the available data for specific waterbody types, the most directly comparable come from the UK Countryside Survey, which looked at trends in headwater stream, ditch and pond plant assemblages across Britain over recent decades. For ponds at least, Countryside Survey results broadly parallel our study findings, showing that in the English lowlands, pond alpha plant richness declined by around 20% (1.8% pa) between 1996 and 2007 (Williams et al 2010). Data from a separate study of ponds in northern England over the same time period show a similar loss of gamma diversity when based on compatible wetland plant species lists (recalculated from Hassall et al., 2012, see Supplementary Information). For English headwater streams, Countryside Survey data showed the reverse trend, with stream plant alpha richness increasing significantly between 1998 and 2007 (Dunbar et al., 2010). This contrasts both with our findings, and with the majority of data from studies of larger European watercourses, which typically show declining alpha and/or gamma plant richness over recent decades (Riis and Sand-Jensen, 2001; Gerhard et al., 2016; Schütz et al., 2008; Steffen, 2013). Comparable data for headwater ditches are hard to come by, with almost all longitudinal studies focused on the wetland flora of more permanent ditches on floodplains, coastal wetlands or semi-natural wetlands, rather than the small, and often seasonal ditches typical of England’s agricultural countryside (Best, 1995; Drake et al., 2010; Whatley, 2013). The main exception is, again, the UK Countryside Survey, which in contrast to our findings, suggests that mean wetland alpha plant richness in headwater ditches increased in the decade before our study was undertaken (Dunbar et al., 2010). Overall, therefore, the results for individual waterbody types provide evidence of a systematic continued loss of plant diversity from agricultural ponds in England’s lowlands, but suggest that trends in stream and ditch flora may be more temporally or spatially variable in agricultural headwaters.

### 4.2 Future losses

Looking forwards, it is clear that the wetland flora in our survey area remains vulnerable to further loss. During the 2010-2018 survey period we directly observed the extinction of species from all catchments and waterbody types through habitat change and destruction including culverting of springs, afforestation of fens and cessation of grazing along waterbody margins. Given that a third (34%) of our catchments’ remaining plant species were restricted to two or fewer sites, and almost a quarter (23%) of these species occurred at exceptionally low abundance, there is a high risk of further loss as a direct result of habitat impacts. An additional threat comes from habitat isolation (Bosiacka et al., 2008). Both theoretical and empirical studies of extinction debt (Tilman et al., 1994; Loehle and Li, 1996) have shown that fragmented habitats with very small populations have high species extinction rates (Halley et al., 2016). Indeed the risk may be especially high for plants located in spatially discrete wetlands, including ponds (Deane et al., 2017; Deane and He, 2018). This suggests a particular vulnerability in our catchments where the majority (over 80%) of plant species with highly restricted distribution occur only in ponds. Ponds also supported the greatest number of rare species in our catchments and, particularly in the Barkby, suffered considerable loss of these taxa during the nine-year study. Extinction debt is held to be especially likely for rare species (Dullinger et al., 2013), which may help to explain why loss of rare species from our catchments has been so high and indicates that the demise of the rare taxa remaining in our catchments may continue.

### 4.3 Biodiversity gain from nature based measures

The addition of nature based measures brought almost immediate biodiversity gains to the two catchments where they were introduced. Creation of clean water ponds, in particular, more than compensated for recent losses and, after five years, had increased whole catchment richness by approximately a quarter and tripled the number of rare plant species present. The rapid colonization rates observed in new nature based ponds is not entirely unexpected. Authors from Darwin onwards have noted the mobility of many freshwater taxa (Darwin, 1859; Talling, 1951; Soons, 2006) whilst other studies have shown that ponds can develop rich plant assemblages within five to six years (Williams, 2010; Mitsch, 2012). New clean water ponds have sometimes been shown to be richer than pre-existing ponds within two to three years (Parikh and Gale, 1998; Williams et al., 2010).

The high proportion of rare species that appeared in the catchment’s clean water ponds is more surprising, but not unprecedented. Williams et al. (1998) showed that 6-12 year-old ponds supported more uncommon plant species than older ponds in lowland Britain, whilst Fleury and Perrin (2004), found that populations of threatened temporary pond plants were greatest in the first 2–3 years after ponds were created. Beyond this, there are many anecdotal examples of nationally rare plant species appearing in newly created ponds, particularly when they are located in semi-natural habitats (e.g. Barnes, 1983; Kennison, 1986; Erskine et al., 2018).

The reason that rare plant species show a propensity to colonise new ponds is not completely clear. Rare species are often held to be poor competitors (Buchele et al.,1992; Shimada and Ishihama, 2000; Lloyd et al., 2002; Cacho and Strauss, 2014) and the bare substrates of new ponds may provide a competitor-free zone in the first few years after creation. Equally, recently excavated waterbodies tend to have relatively nutrient-poor substrates which may directly benefit groups including charophytes that have been shown to thrive below relatively low nutrient thresholds (Lambert and Davy, 2011). Alternatively, remnants of soil left after pond creation may simply have exposed previously buried seed banks, allowing rare plants that are no longer present in the standing flora to germinate (Nishihiro et al., 2005). In our study, it seems likely that more than one factor was at play. A proportion of the nationally and regionally rare species that we recorded (e.g. *Triglochin palustris, Juncus subnodulosus, Isolepis setacea*) appeared only in ponds created in an area of secondary woodland partly planted on a former fen. *Triglochin palustris*, at least, had previously been recorded from the fen, and although our pre-excavation surveys did not find it in the area where the ponds were created, there seems a high likelihood that this species germinated from a pre-existing seed bank. Other taxa, including *Hippuris vulgaris* and *Chara* species, appeared in new ponds that were located in isolated dry ground areas where colonising species can only have arrived though wind or bird transported propagules (Soons, 2006; Soonsl et al., 2008). Here, their successful colonisation presumably reflects the new opportunities provided by the ponds’ bare substrates.

It is interesting to note that our data provide circumstantial evidence that adding new ponds helped to increase the biodiversity of pre-existing waterbodies. Of the three catchments, the Stonton, which received most nature based measures, also saw the lowest rate of species loss. Indeed, the only pre-existing waterbody type to show a net increase in total richness during the survey period were the Stonton ponds; the habitat type which would be expected to benefit most from the increased connectivity and dispersal opportunities created by adding new ponds in their vicinity. Looking in detail at species trends provides evidence to support this. For example, the submerged aquatic *Potamogeton pusillus*, was not present in the Stonton catchment before the nature-based measures were introduced. However, it rapidly colonised a number of the new clean water ponds, and was subsequently recorded in first one, and then a second pre-existing pond, around 1 km distant. Such incidents provide a strong indication that in some cases, the new ponds have acted as stepping stones, which ultimately helped to support greater alpha and gamma richness in the Stonton’s pre-existing waterbodies.

### 4.4 Effectiveness of ecosystem services waterbodies

There is often an assumption that nature based measures will be ‘good for wildlife’, and that ecosystem services features, such as SUDs waterbodies, agricultural bunded ditches and interception ponds will inevitably provide multi-functional benefits that include biodiversity gain (Burgess-Gamble et al., 2017). In practice, studies to test these assumptions have shown mixed results depending on the type of measures introduced and the biotic group used to derive metrics (Hansson et al., 2005; Thiere et al., 2009; Wiegleb et al., 2017). The results from our plant-based study provides some support for the value of ecosystem services waterbodies, particularly for reversing the impact of short term losses in catchment plant richness, and increasing the number of populations of some unique and replicate species. However, ecosystem services waterbodies did not restore the loss of rare plant species from our catchments.. These findings tally with the majority of other plant-based studies of waterbodies and wetlands which have shown that ecosystem service features tend to support more homogeneous communities, and fewer high quality or rare plant species, than their semi-natural equivalents (Balcombe et al., 2005; Brooks et al., 2005; Aronson and Galatowitsch, 2008; Robertson et al., 2018; Price et al., 2019). Taken together, these findings suggest that ecosystem service measures may have a role in helping to support catchment biodiversity. However, it is important not to inflate their potential value beyond the level justified by evidence.

### 4.5 Will the gains persist?

Whether the catchments’ biodiversity gains will persist in the longer-term remains an open question. Evidence from other new clean water ponds shows that this waterbody type can retain high biodiversity for many decades (Williams et al., 2007, Williams et al., 2010). However, in the majority of cases these ponds were created in semi-natural landscapes. Further monitoring is required to determine whether the clean water ponds in our agricultural landscapes can maintain their disproportionate contribution to catchment biodiversity. The future biodiversity benefits from the ecosystem services ponds seems more doubtful. Evidence suggests that new ponds in highly impacted landscapes tend to reach their greatest richness within the first five years after creation and then decline. This is almost certainly because they degrade once they begin to fill with polluted sediment and water (Williams, 2007 Robertson et al., 2018). For on-line ecosystem service features like ours, where the waterbody’s main function is to intercept polluted sediment and water, short to medium term declines in biodiversity value seem likely. Some decline is already evident from our data: a number of ecosystem services waterbodies supported uncommon plants (*Potamogeton* and *Chara species)* in the first few years after creation which were rapidly lost as these features began to fill with sediment and became more algal dominated. It is possible that regularly desilting ecosystem services waterbodies could remove sediment and pollutants and return them to an earlier, botanically richer successional stage, but this possibility is yet to be tested. Further surveillance data are essential to measure both the extent of any further losses, and the potential for de-silting to reset the clock.

## 5 Implications

### 5.1 The value of gamma richness data

Alpha richness measures remain a mainstay of ecological research. However, over the last few decades there has been an increasing trend towards collection of gamma data, particularly for freshwater habitats (Bubíková and Hrivnák, 2018). Most studies measure gamma richness as summed alpha values rather than adopting what can sometimes be an impossibly time consuming census approach. In this study we used wetland plants which are a relatively quick and easy group to survey in order to undertake census surveys of waterbodies, and found that this provided advantages for assessing catchment-scale change. For example, alpha data could not provide evidence that the observed species losses were real because between-year variability in site richness confounded statistical significance tests. Gamma census data, which reveal the real world, not only made it possible to measure losses and gains over the relatively short time-scale of the project, but provided confidence in measuring attributes such as the number of restricted species, area of occupancy, and species movement between waterbodies. We suggest that census data could be useful for other freshwater studies, particularly to identify trends over short timescales where changes are likely to be subtle. Studies that investigate the net change across a wider range of landscapes (agricultural, urban or semi-natural) would be particularly welcome in order to place the results recorded here in a wider context.

### 5.2 Ponds as catchment controlling habitats

Ponds were a key habitat in the agricultural catchments that we studied. By a considerable margin, ponds supported the greatest number of freshwater plant species, the most uncommon species and the highest proportion of unique taxa, with 40% of species only found in ponds. Loss of pond species, particularly submerged aquatic taxa, had a disproportionately high impact on catchment richness and rarity trends.

Recognition of the importance of ponds for supporting catchment biodiversity has been growing for almost two decades (reviewed in Biggs et al., 2017 and Hill et al., 2018). Calls for greater representation of ponds within national and international policy have been growing apace (Williams et al., 2004; Kristensen and Globevnik, 2014; Sayer, 2014; Hassall, 2014; Biggs et al., 2017). Yet there is still no requirement to monitor ponds as part of water quality assessment in Europe, the US or other countries where river monitoring is mandatory (Hill et al., 2018). Equally lacking are water and nature conservation policies that support pond protection at site or catchment level (e.g. River Basin Management Plans) despite increasing evidence that, *en masse*, small habitats like ponds are critical for maintaining landscape scale biodiversity (Hill et al., 2018; Grasel et al., 2018; Fahrig, 2019). The evidence from our study adds further weight to the argument that ponds need to be specifically included in policy and legislation if we are to reduce freshwater biodiversity loss and stand a chance of making sustainable catchment management a reality.

### 5.3 Good news for biodiversity protection

Amidst the gloomy reality of global declines in freshwater habitats and species, the results from this study provide evidence that proactive habitat creation measures can make a positive difference to agricultural catchment-scale biodiversity over comparatively short time scales. Our results provide the first demonstration of a whole-landscape increase in freshwater biodiversity as a result of agricultural land management measures. The findings emphasise the potential for clean water pond creation to bring significant benefits to catchment biodiversity in agricultural landscapes.

## 6 Acknowledgements

This work was supported by the Environment Agency (Grants CRF013 and ENVREAN002188) and by Syngenta Ltd who funded a Post-Doctoral Research Assistant post as part of the project from 2010-15. We are enormously grateful to the landowners and tenants in all catchments for granting access to sampling sites, and particularly to landowners in the Eye and Stonton catchments for their close involvement with the introduction of measures for the project. Thanks to Sian Davies, Anita Casey, Adrianna Hawczak and Elaine McGoff for their practical and technical support. We are also pleased to acknowledge the advice and guidance provided by Lorriane Maltby, the Welland Valley Partnership and Welland Rivers Trust, and to Bill Brierley who helped initiate the project. We are very grateful to the anonymous referees who’s comments allowed us to substantially improve this manuscript.

## References

Aalto, E.A., Micheli, F., Boch, C.A., Espinoza Montes, J.A., Woodson, C.B. and De Leo, G.A., 2019. Catastrophic Mortality, Allee Effects, and Marine Protected Areas. The American Naturalist, 193 (3), pp.391–408. https://doi.org/10.1086/701781

Aronson, M.F. and Galatowitsch, S., 2008. Long-term vegetation development of restored prairie pothole wetlands. Wetlands, 28(4), pp.883–895. https://doi.org/10.1672/08-142.1

Balcombe, C.K., Anderson, J.T., Fortney, R.H., Rentch, J.S., Grafton, W.N. and Kordek, W.S., 2005. A comparison of plant communities in mitigation and reference wetlands in the mid-Appalachians. Wetlands, 25 (1), pp.130–142. 10.1672/0277-5212(2005)025[0130:ACOPCI]2.0.CO;2

Barnes, L.E., 1983. The colonization of ball-clay ponds by macroinvertebrates and macrophytes. Freshwater Biology, 13 (6), pp.561–578. https://doi.org/10.1111/j.1365-2427.1983.tb00013.x

Bernhardt, E.S. and Palmer, M.A., 2011. River restoration: the fuzzy logic of repairing reaches to reverse catchment scale degradation. Ecological applications, 21(6), pp.1926–1931. https://doi.org/10.1890/10-1574.1

Best, E.P.H., Van der Schaaf, S. and Oomes, M.J.M., 1995. Responses of restored grassland ditch vegetation to hydrological changes, 1989–1992. Vegetatio, 116 (2), pp.107–122. https://doi.org/10.1007/BF00045302

Biggs, J., Stoate, S., Williams, P., Brown, C., Casey, A., Davies, S., Diego, I.G., Hawczak, A., Kizuka, T., McGoff, E. and Szczur, J., 2014. Water Friendly Farming: Results and practical implications of the first 3 years of the programme. Freshwater Habitats Trust, Oxford, and Game and name>Wildlife Conservation Trust, Fordingbridge, UK.

Biggs, J., Von Fumetti, S. and Kelly-Quinn, M., 2017. The importance of small waterbodies for biodiversity and ecosystem services: implications for policy makers. Hydrobiologia, 793 (1), pp.3–39. https://doi.org/10.1007/s10750-016-3007-0

Bosiacka, B., Pacewicz, K. and Pienkowski, P., 2008. Spatial analysis of plant species distribution among small water bodies in a agricultural landscape. Acta agrobotanica, 61 (2).

Brooks, R.P., Wardrop, D.H., Cole, C.A. and Campbell, D.A., 2005. Are we purveyors of wetland homogeneity?: A model of degradation and restoration to improve wetland mitigation performance. Ecological Engineering, 24(4), pp.331–340. https://doi.org/10.1016/j.ecoleng.2004.07.009

Brown, C.D., Turner, N., Hollis, J., Bellamy, P., Biggs, J., Williams, P., Arnold, D., Pepper, T. and Maund, S., 2006. Morphological and physico-chemical properties of British aquatic habitats potentially exposed to pesticides. Agriculture, ecosystems & environment, 113(1-4), pp.307–319. https://doi.org/10.1016/j.agee.2005.10.015

BSBI maps. 2019. Species recorded during the period 2000 to 2018. https://bsbi.org/maps. Accessed 15th February 2019.

Bubíková, K. and Hrivnák, R., 2018. Comparative macrophyte diversity of waterbodies in the Central European landscape. Wetlands, 38(3), pp.451–459. https://doi.org/10.1007/s13157-017-0987-0

Buchele, D.E., Baskin, J.M. and Baskin, C.C., 1992. Ecology of the endangered species Solidago shortii. V. Plant associates. Bulletin of the Torrey Botanical Club, pp.208–213. https://doi.org/10.2307/2997032

Burgess-Gamble, L., Ngai, R., Wilkinson, M., Nisbet, T., Pontee, N., Harvey, R., Kipling, K., Addy, S., Rose, S., Maslen, S. and Jay, H., 2017. Working with Natural Processes–Evidence Directory. Environmental Agency, Report No. SC150005.

Cacho, N.I. and Strauss, S.Y., 2014. Occupation of bare habitats, an evolutionary precursor to soil specialization in plants. Proceedings of the National Academy of Sciences, 111 (42.), pp.15132–15137. https://doi.org/10.1073/pnas.1409242111

Caissie, D., 2006. The thermal regime of rivers: a review. Freshwater biology, 51 (8), pp.1389–1406. https://doi.org/10.1111/j.1365-2427.2006.01597.x

Carvalho, L., Mackay, E.B., Cardoso, A.C., Baattrup-Pedersen, A., Birk, S., Blackstock, K.L., Borics, G., Borja, A., Feld, C.K., Ferreira, M.T. and Globevnik, L., 2019. Protecting and restoring Europe’s waters: An analysis of the future development needs of the Water Framework Directive. Science of the Total Environment, 658, pp.1228–1238. https://doi.org/10.1016/j.scitotenv.2018.12.255

Céréghino, R., Biggs, J., Oertli, B. and Declerck, S., 2007. The ecology of European ponds: defining the characteristics of a neglected freshwater habitat. In Pond Conservation in Europe (pp. 1–6). Springer, Dordrecht. https://doi.org/10.1007/s10750-007-9225-8

Cohen-Shacham, E., Walters, G., Janzen, C. and Maginnis, S., 2016. Nature-based solutions to address global societal challenges. IUCN, Gland, Switzerland, 97. https://doi.org/10.2305/IUCN.CH.2016.13.

enColwell, R.K., Chao, A., Gotelli, N.J., Lin, S.Y., Mao, C.X., Chazdon, R.L. and Longino, J.T., 2012. Models and estimators linking individual-based and sample-based rarefaction, extrapolation and comparison of assemblages. Journal of plant ecology, 5 (1), pp.3–21. https://doi.org/10.1093/jpe/rtr044

Cuttle, S.P., Newell-Price, J.P., Harris, D., Chadwick, D.R., Shepherd, M.A., Anthony, S.G.A., Macleod, C.J.A., Haygarth, P.M. and Chambers, B.J., 2016. A method-centric ‘User Manual’for the mitigation of diffuse water pollution from agriculture. Soil Use and Management, 32, pp.162–171. https://doi.org/10.1111/sum.12242

Darwin, C., 1859. The Origin of Species; And, the Descent of Man. Modern library.

Davies, B., Biggs, J., Williams, P., Whitfield, M., Nicolet, P., Sear, D., Bray, S. and Maund, S., 2008. Comparative biodiversity of aquatic habitats in the European agricultural landscape. Agriculture, Ecosystems & Environment, 125 (1-4), pp.1–8. https://doi.org/10.1016/j.agee.2007.10.006

Deane, D.C. and He, F., 2018. Loss of only the smallest patches will reduce species diversity in most discrete habitat networks. Global change biology, 24 (12), pp.5802–5814. https://doi.org/10.1111/gcb.14452

Deane, D.C., Fordham, D.A., He, F. and Bradshaw, C.J., 2017. Future extinction risk of wetland plants is higher from individual patch loss than total area reduction. Biological Conservation, 209, pp.27–33. https://doi.org/10.1016/j.biocon.2017.02.005

Defra, 2018. The guide to cross compliance in England 2018. https://assets.publishing.service.gov.uk/government/uploads/system/uploads/attachment_data/file/668684/Cross_Compliance_2018_guide_v1.0.pdf (accessed 13 April 2019)

Drake, C.M., Stewart, N.F., Palmer, M.A. and Kindemba, V.L., 2010. The ecological status of ditch systems: an investigation into the current status of the aquatic invertebrate and plant communities of grazing marsh ditch systems in England and Wales. Buglife-The Invertebrate Conservation Trust: Peterborough, UK.

Duffy, J.E., 2009. Why biodiversity is important to the functioning of real-world ecosystems. Frontiers in Ecology and the Environment, 7 (8), pp.437–444. https://doi.org/10.1890/070195

Dullinger, S., Essl, F., Rabitsch, W., Erb, K.H., Gingrich, S., Haberl, H., Hülber, K., Jarošík, V., Krausmann, F., Kühn, I. and Pergl, J., 2013. Europe’s other debt crisis caused by the long legacy of future extinctions. Proceedings of the National Academy of Sciences, 110 (18), pp.7342–7347. https://doi.org/10.1073/pnas.1216303110

Dunbar, M., Murphy, J., Clarke, R., Baker, R., Davies, C., Scarlett, P. 2010 Countryside Survey: Headwater Streams Report from 2007. Technical Report No. 8/07 NERC/Centre for Ecology & Hydrology pp 67.

Erskine, S.E., Killick, H.J., Lambrick C.R. and Lee E.M., 2018. Oxfordshire’s Threatened Plants. Pisces Publications.

Fahrig, L., 2019. Habitat fragmentation: A long and tangled tale. Global Ecology and Biogeography, 28 (1), pp.33–41. https://doi.org/10.1111/geb.12839

Fleury, Z. and Perrin, C.S., 2004. Vegetation colonisation of temporary ponds newly dug in the marshes of the Grande Caricaie (lake of Neuchatel, Switzerland). Archives des Sciences, 57 (2–3), pp.105–112.

Foley, J.A., DeFries, R., Asner, G.P., Barford, C., Bonan, G., Carpenter, S.R., Chapin, F.S., Coe, M.T., Daily, G.C., Gibbs, H.K. and Helkowski, J.H., 2005. Global consequences of land use. science, 309 (5734), pp.570–574. https://doi.org/10.1126/science.1111772

Freshwater Habitats Trust, 2015. Wetland plant species list. https://freshwaterhabitats.org.uk/wp-content/uploads/2015/03/35-WETLAND-PLANTS-LATIN-RECORDING-FORM-FINAL.pdf (accessed 10 December 2018)

Freshwater Habitats Trust, 2019. Pond Creation Toolkit https://freshwaterhabitats.org.uk/projects/million-ponds/pond-creation-toolkit/ (accessed 20 March 2019).

Gołdyn, H., 2010. Changes in plant species diversity of aquatic ecosystems in the agricultural landscape in West Poland in the last 30 years. Biodiversity and conservation, 19 (1), pp.61–80. https://doi.org/10.1007/s10531-009-9702-7

Gordon, L.J., Peterson, G.D. and Bennett, E.M., 2008. Agricultural modifications of hydrological flows create ecological surprises. Trends in ecology & evolution, 23 (4), pp.211–219. https://doi.org/10.1016/j.tree.2007.11.011

Gotelli, N.J. and Colwell, R.K., 2011. Estimating species richness. Biological diversity: frontiers in measurement and assessment, 12, pp.39–54.

Grasel, D., Mormul, R.P., Bozelli, R.L., Thomaz, S.M. and Jarenkow, J.A., 2018. Brazil’s Native Vegetation Protection Law threatens to collapse pond functions. Perspectives in ecology and conservation, 16(4), pp.234–237. https://doi.org/10.1016/j.pecon.2018.08.003

Halley, J.M., Monokrousos, N., Mazaris, A.D., Newmark, W.D. and Vokou, D., 2016. Dynamics of extinction debt across five taxonomic groups. Nature communications, 7, p.12283. https://doi.org/10.1038/ncomms12283

Hansson, L.A., Brönmark, C., Anders Nilsson, P. and Åbjörnsson, K., 2005. Conflicting demands on wetland ecosystem services: nutrient retention, biodiversity or both?. Freshwater Biology, 50 (4), pp.705–714. https://doi.org/10.1111/j.1365-2427.2005.01352.x

Harris, G.P. and Heathwaite, A.L., 2012. Why is achieving good ecological outcomes in rivers so difficult?. Freshwater Biology, 57, pp.91–107. https://doi.org/10.1111/j.1365-2427.2011.02640.x

Hassall, C., 2014. The ecology and biodiversity of urban ponds. Wiley Interdisciplinary Reviews: Water, 1 (2), pp.187–206. https://doi.org/10.1002/wat2.1014

Hassall, C., Hollinshead, J. and Hull, A., 2012. Temporal dynamics of aquatic communities and implications for pond conservation. Biodiversity and Conservation, 21(3), pp.829–852. https://doi.org/10.1007/s10531-011-0223-9

Hill, J.E., 2016. Collapse of a reproducing population of non-native African jewelfish (Hemichromis letourneuxi) in a Florida lake. NeoBiota, 29, p.35. https://doi.org/10.3897/neobiota.29.7213

Hill, M.J., Hassall, C., Oertli, B., Fahrig, L., Robson, B.J., Biggs, J., Samways, M.J., Usio, N., Takamura, N., Krishnaswamy, J. and Wood, P.J., 2018. New policy directions for global pond conservation. Conservation Letters, 11 (5), p.e12447. https://doi.org/10.1111/conl.12447

Hisano, M., Searle, E.B. and Chen, H.Y., 2018. Biodiversity as a solution to mitigate climate change impacts on the functioning of forest ecosystems. Biological Reviews, 93 (1), pp.439–456. https://doi.org/10.1111/brv.12351

JNCC, Conservation designations spreadsheet. Last updated July 2018. http://jncc.defra.gov.uk/page-3408. Accessed 15th February 2019

IPBES. 2019. Summary for policymakers of the global assessment report on biodiversity and ecosystem services of the Intergovernmental Science-Policy Platform on Biodiversity and Ecosystem Services. S. Díaz, J. Settele, E. S. Brondizio E.S., H. T. Ngo, M. Guèze, J. Agard, A. Arneth, P. Balvanera, K. A. Brauman, S. H. M. Butchart, K. M. A. Chan, L. A. Garibaldi, K. Ichii, J. Liu, S. M. Subramanian, G. F. Midgley, P. Miloslavich, Z. Molnár, D. Obura, A. Pfaff, S. Polasky, A. Purvis, J. Razzaque, B. Reyers, R. Roy Chowdhury, Y. J. Shin, I. J. Visseren-Hamakers, K. J. Willis, and C. N. Zayas (eds.). IPBES secretariat, Bonn, Germany. https://www.ipbes.net/global-assessment-report-biodiversity-ecosystem-services Accessed 24th October 2019

Kennison, G.C.B., 1986. Preliminary observations on the plant colonisation of experimental turf ponds in a Broadland fen. Transactions of the Norfolk Naturalists Society, 27, pp.193–198.

Kristensen, P. and Globevnik, L., 2014. European small water bodies. In Biology and Environment: Proceedings of the Royal Irish Academy (Vol. 114, No. 3, pp. 281–287). Royal Irish Academy. https://doi.org/10.3318/BIOE.2014.13

Lambert, S.J. and Davy, A.J., 2011. Water quality as a threat to aquatic plants: discriminating between the effects of nitrate, phosphate, boron and heavy metals on charophytes. New Phytologist, 189(4), pp.1051–1059. https://doi.org/10.1111/j.1469-8137.2010.03543.x

Lloyd, K.M., Lee, W.G. and Wilson, J.B., 2002. Competitive abilities of rare and common plants: comparisons using Acaena (Rosaceae) and Cnionocnloa (Poaceae) from New Zealand. Conservation Biology, 16 (4), pp.975–985. https://doi.org/10.1046/j.1523-1739.2002.01033.x

Law, A., Baker, A., Sayer, C., Foster, G., Gunn, I.,Taylor, P., Pattison, Z.., Blaikie, J., Willby, N., In press. The effectiveness of aquatic plants as surrogates for wider biodiversity in standing fresh waters. Freshwater Biology (manuscript ID FWB-P-Mar-19-0124.R1)

Loehle, C. and Li, B.L., 1996. Habitat destruction and the extinction debt revisited. Ecological applications, 6 (3), pp.784–789. https://doi.org/10.2307/2269483

Louhi, P., Mykrä, H., Paavola, R., Huusko, A., Vehanen, T., Mäki-Petäys, A. and Muotka, T., 2011. Twenty years of stream restoration in Finland: little response by benthic macroinvertebrate communities. Ecological Applications, 21 (6), pp.1950–1961. https://doi.org/10.1890/10-0591.1

Mariotte, P., Vandenberghe, C., Kardol, P., Hagedorn, F. and Buttler, A., 2013. Subordinate plant species enhance community resistance against drought in semi-natural grasslands. Journal of Ecology, 101 (3), pp.763–773. https://doi.org/10.1111/1365-2745.12064

Mitsch, W.J., Zhang, L., Stefanik, K.C., Nahlik, A.M., Anderson, C.J., Bernal, B., Hernandez, M. and Song, K., 2012. Creating wetlands: primary succession, water quality changes, and self-design over 15 years. Bioscience, 62 (3), pp.237–250. https://doi.org/10.1525/bio.2012.62.3.5

Moss, B., 2007. Water pollution by agriculture. Philosophical Transactions of the Royal Society B: Biological Sciences, 363 (1491), pp.659–666. https://doi.org/10.1098/rstb.2007.2176

Nishihiro, J., Nishihiro, M.A. and Washitani, I., 2006. Assessing the potential for recovery of lakeshore vegetation: species richness of sediment propagule banks. Ecological Research, 21(3), pp.436–445. https://doi.org/10.1007/s11284-005-0133-y

Oertli, B., 2018. Freshwater biodiversity conservation: The role of artificial ponds in the 21st century. Aquatic Conservation: Marine and Freshwater Ecosystems, 28(2), pp.264–269. https://doi.org/10.1002/aqc.2902

Palmer, M.A., 2009. Reforming watershed restoration: science in need of application and applications in need of science. Estuaries and coasts, 32 (1), pp.1–17. https://doi.org/10.1007/s12237-008-9129-5

Parikh, A. and Gale, N., 1998. Vegetation monitoring of created dune swale wetlands, Vandenberg Air Force Base, California. Restoration Ecology, 6 (1), pp.83–93. https://doi.org/10.1046/j.1526-100x.1998.06111.x

Price, E.P., Spyreas, G. and Matthews, J.W., 2018. Biotic homogenization of regional wetland plant communities within short time-scales in the presence of an aggressive invader. Journal of Ecology, 106(3), pp.1180–1190. https://doi.org/10.1002/eap.1827

Rayment, M., 2017. Assessing the costs of Environmental Land Management in the UK.

Rhodes, H.M., Closs, G.P. and Townsend, C.R., 2007. Stream ecosystem health outcomes of providing information to farmers and adoption of best management practices. Journal of applied ecology, 44(6), pp.1106–1115. https://doi.org/10.1111/j.1365-2664.2007.01397.x

Riis, T. and Sand-Jensen, K., 2001. Historical changes in species composition and richness accompanying perturbation and eutrophication of Danish lowland streams over 100 years. Freshwater Biology, 46(2), pp.269–280. https://doi.org/10.1046/j.1365-2427.2001.00656.x.

Robertson, M., Galatowitsch, S.M. and Matthews, J.W., 2018. Longitudinal evaluation of vegetation richness and cover at wetland compensation sites: implications for regulatory monitoring under the Clean Water Act. Wetlands Ecology and Management, 26(6), pp.1089–1105. https://doi.org/10.1007/s11273-018-9633-8

Roni, P., Beechie, T., Pess, G. and Hanson, K., 2014. Wood placement in river restoration: fact, fiction, and future direction. Canadian Journal of Fisheries and Aquatic Sciences, 72(3), pp.466–478. https://doi.org/10.1139/cjfas-2014-0344

Sayer, C.D., 2014. Conservation of aquatic landscapes: ponds, lakes, and rivers as integrated systems. Wiley Interdisciplinary Reviews: Water, 1 (6), pp.573–585. https://doi.org/10.1002/wat2.1045

Schütz, W., Veit, U. and Kohler, A., 2008. The aquatic vegetation of the Upper Danube river-past and present. Large Rivers, pp.167–191. https://doi.org/10.1127/lr/18/2008/167

Shimada, M. and Ishihama, F., 2000. Asynchronization of local population dynamics and persistence of a metapopulation: a lesson from an endangered composite plant, Aster kantoensis. Population Ecology, 42 (1), pp.63–72. https://doi.org/10.1007/s101440050010

Soons, M.B., 2006. Wind dispersal in freshwater wetlands: knowledge for conservation and restoration. Applied Vegetation Science, 9 (2), pp.271–278. https://doi.org/10.1111/j.1654-109X.2006.tb00676.x

Soons, M.B., Van Der Vlugt, C., Van Lith, B., Heil, G.W. and Klaassen, M., 2008. Small seed size increases the potential for dispersal of wetland plants by ducks. Journal of Ecology, 96 (4), pp.619–627. https://doi.org/10.1111/j.1365-2745.2008.01372.x

Steffen, K., Becker, T., Herr, W. and Leuschner, C., 2013. Diversity loss in the macrophyte vegetation of northwest German streams and rivers between the 1950s and 2010. Hydrobiologia, 713 (1), pp.1–17. https://doi.org/10.1007/s10750-013-1472-2

Stroh, P., Leach, S.J., August, T.A., Walker, K.J., Pearman, D.A., Rumsey, F.J., Harrower, C.A., Fay, M.F., Martin, J.P., Pankhurst, T., Preston, C.D., Taylor, I., 2014. A Vascular Plant Red List for England. Botanical Society of Britain & Ireland.

Stubbington, R., 2012. The hyporheic zone as an invertebrate refuge: a review of variability in space, time, taxa and behaviour. Marine and Freshwater Research, 63(4), pp.293–311. https://doi.org/10.1071/MF11196

Talling, J.F., 1951. The element of chance in pond populations. The Naturalist, 1951, pp.157–170.

Thiere, G., Milenkovski, S., Lindgren, P.E., Sahlén, G., Berglund, O. and Weisner, S.E., 2009. Wetland creation in agricultural landscapes: biodiversity benefits on local and regional scales. Biological conservation, 142 (5), pp.964–973. https://doi.org/10.1016/j.biocon.2009.01.006

Thomas, R.E., Teel, T., Bruyere, B. and Laurence, S., 2018. Metrics and outcomes of conservation education: a quarter century of lessons learned. Environmental Education Research, pp.1–21. https://doi.org/10.1080/13504622.2018.1450849

Tilman, D., May, R.M., Lehman, C.L. and Nowak, M.A., 1994. Habitat destruction and the extinction debt. Nature, 371 (6492), p.65. https://doi.org/10.1038/371065a0

Whatley, M.H., van Loon, E.E., van Dam, H., Vonk, J.A., van der Geest, H.G. and Admiraal, W., 2014. Macrophyte loss drives decadal change in benthic invertebrates in peatland drainage ditches. Freshwater Biology, 59 (1), pp.114–126. https://doi.org/10.1111/fwb.12252

Wiegleb, G., Dahms, H.U., Byeon, W.I. and Choi, G., 2017. To what extent can constructed wetlands enhance biodiversity. International Journal of Environmental Science and Development, 8 (8), pp.561–569. https://doi.org/10.18178/ijesd.2017.8.8.1016

Wiegleb, G., Gebler, D., van de Weyer, K. and Birk, S., 2016. Comparative test of ecological assessment methods of lowland streams based on long-term monitoring data of macrophytes. Science of the Total Environment, 541, pp.1269–1281. https://doi.org/10.1016/j.scitotenv.2015.10.005

Williams, P., Biggs, J., Crowe, A., Murphy, J., Nicolet, P., Weatherby, A., Dunbar, M., 2010. Countryside Survey: Ponds Report from 2007. Technical Report No.7/07. Pond Conservation and NERC/Centre for Ecology & Hydrology..

Williams, P., Whitfield, M. and Biggs, J., 2007. How can we make new ponds biodiverse? A case study monitored over 7 years. In Pond Conservation in Europe (pp. 137–148). Springer, Dordrecht. https://doi.org/10.1007/978-90-481-9088-1_12

Williams, P., Whitfield, M., Biggs, J., Bray, S., Fox, G., Nicolet, P. and Sear, D., 2004. Comparative biodiversity of rivers, streams, ditches and ponds in an agricultural landscape in Southern England. Biological conservation, 115(2), pp.329–341. https://doi.org/10.1016/S0006-3207(03)00153-8

Williams, P.J., Biggs J, Barr, C.J., Cummins, C.P., Gillespie, M.K., Rich, T.C.G., Baker, A., Beesley, J., Corfield, A., Dobson, D., Culling, A.S., Fox, G., Howard, D.C., Luursema, K., Rich, M., Samson, D., Scott, W.A., White, R. and Whitfield, M., 1998. Lowland Pond Survey 1996. Department of the Environment, Transport and the Regions.

Zelnik, I., Gregorič, N. and Tratnik, A., 2018. Diversity of macroinvertebrates positively correlates with diversity of macrophytes in karst ponds. Ecological Engineering, 117, pp.96–103.

Zhang, Y., Collins, A.L., Jones, J.I., Johnes, P.J., Inman, A. and Freer, J.E., 2017. The potential benefits of on-farm mitigation scenarios for reducing multiple pollutant loadings in prioritised agri-environment areas across England. Environmental Science & Policy, 73, pp.100–114. https://doi.org/10.1016/j.envsci.2017.04.004

